# A statistical framework for measuring reproducibility and replicability of high-throughput experiments from multiple sources

**DOI:** 10.1101/827527

**Authors:** Monia Ranalli, Yafei Lyu, Hillary Koch, Qunhua Li

## Abstract

Replication is essential to reliable and consistent scientific discovery in high-throughput experiments. Quantifying the replicability of scientific discoveries and identifying sources of irreproducibility have become important tasks for quality control and data integration. In this work we introduce a novel statistical model to measure the reproducibility and replicability of findings from replicate experiments in multi-source studies. Using a nested copula mixture model that characterizes the interdependence between replication experiments both across and within sources, our method quantifies reproducibility and replicability of each candidate simultaneously in a coherent framework. Through simulation studies, an ENCODE ChIP-seq dataset and a SEQC RNA-seq dataset, we demonstrate the effectiveness of our method in diagnosing the source of discordance and improving the reliability of scientific discoveries.

## 1. Introduction

High-throughput technologies play an important role in modern biological studies. By profiling a large number of candidates in a single experiment, high-throughput technologies enable researchers to efficiently identify potential targets for downstream analyses. In the past decade, thousands of high-throughput genomics datasets have been produced by individual labs and multi-institutional consortium studies, such as ENCODE (ENCODE Consortium et al., 2012) and Roadmap projects (Roadmap Epigenomics Consortium et al., 2015). Many datasets measure the same biological traits, but are generated from different labs or different studies. Integration of these data can potentially enhance statistical power and improve the reliability of scientific interpretation. However, data are affected by source-specific factors, such as laboratory conditions, reagent lots, and personnel difference (Leek *and others*, 2010). Variation in such factors can create inconsistent measurements across sources even if the underlying biology is identical, compromising the reliability of downstream analyses and possibly leading to incorrect conclusions.

Two terms are often used to describe the consistency of scientific discoveries in multi-source studies: replicability and reproducibility (Patil *and others*, 2016; Plesser, 2018; Leek and Peng, 2015; National Academies of Sciences and *others*, 2016). The definitions of these terms are often conflated in literature and sometimes variable depending on the context and the setup of replications (Plesser, 2018; National Academies of Sciences *and others*, 2016). In high-throughput multi-source studies, the relationship among replication experiments can be described as a nested structure (Figure 1): replicate experiments from the same source are usually performed using the same experimental and data-analytical workflow, and those from different sources are performed independently using possibly different workflows. The measurements from the replicates from the same source are usually more consistent to each other than measurements from the replicates across different sources. To fit this nested structure, we adapted the statistical definition for reproducibility and replicability in Patil *and others* (2016) as follows: Replicability assesses the consistency between findings from replicate samples measured by *different* labs; whereas reproducibility assesses the consistency between findings from replicate samples measured by the *same* lab. Each term reflects different sources of heterogeneity: the former generally reflects the difficulty of replication, which may be related to the nature of the biology or the technology; whereas the latter mainly reflects the within-lab variability in measurement and is often related to lab-specific factors. Dissecting the discordance across experiments in terms of reproducibility and replicability will not only help investigators to understand sources of variation and identify actions for improving the reliability of the results, but also help take proper account of the heterogeneity within and across sources in data integration.

**Fig. 1.**
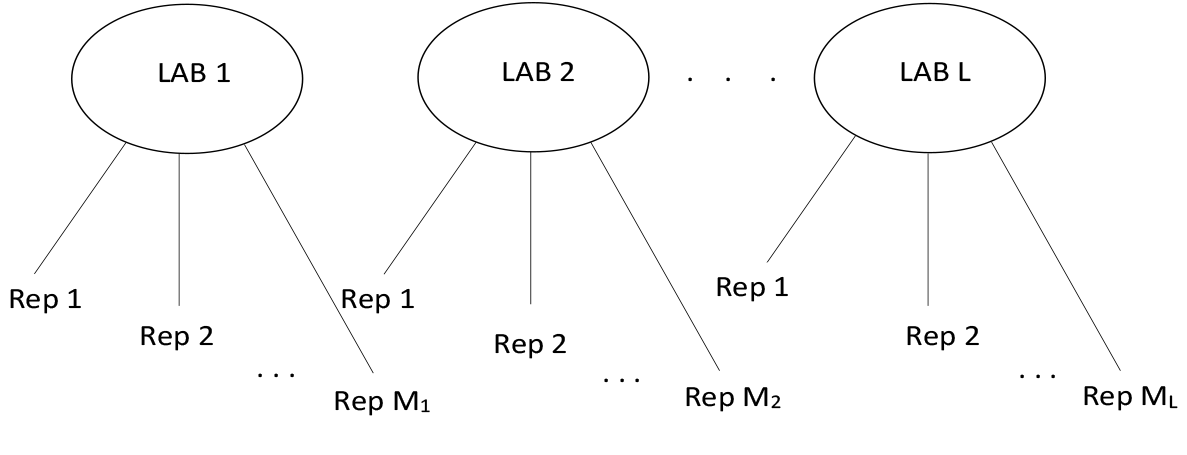
Typical experimental design in consortium studies.

The primary output of a high-throughput experiment is typically organized into a ranked candidate list, on which the candidates are ranked by the strength of their association to the biological trait of interest. For example, the candidate list for a ChIP-seq experiment of a transcription factor consists of the putative protein binding sites on the DNA, along with the p-values that these genomic loci are bound by the transcription factor. A traditional way to measure the consistency between the ranked candidate lists from two datasets is to compute Spearman’s rank correlation coefficient between the two lists. Although this method reflects the overall concordance of two candidate lists, it does not quantify the concordance of individual candidates across datasets. To address this issue, Li *and others* (2011) proposed a copula mixture model. This method assesses the reproducibility of each candidate on the candidate lists based on a pair of replicate samples and reports the expected rate of irreproducible discoveries in the selected candidates, termed the irreproducible discovery rate (IDR), in a manner analogous to the false discovery rate (FDR). The IDR method and its variants have been used for assessing the reproducibility of ChIP-seq studies (Landt *and others*, 2012; Bailey *and others*, 2013) and identifying differentially expressed genes that are concordant across microarray and RNA-seq experiments from the same biological sample (Lyu and Li, 2016).

However, although the IDR model is appropriate for assessing the reproducibility of replicate experiments from one source, it is not entirely suited for evaluating consistency of findings from different sources. This is because it is source-agnostic; that is, it treats replicate samples without regard to their sources. Consequently, within-source dependence induced by source-specific variation are not accounted for. Moreover, it does not distinguish between reproducibility and replicability, and cannot provide information about the sources of heterogeneity.

Here we present nestedIDR, a novel nested mixture copula model to simultaneously measure the reproducibility and replicability of findings from high-throughput experiments. By extending the IDR method to model the nested structure inherent to replicate samples within and across sources, this method effectively takes account of source-specific heterogeneity. As we will show, it not only allows one to track down the sources of discordance across replicate experiments, but also improves the power and robustness to identify biologically relevant candidates.

Section 2 introduces the nested mixture copula model and its estimation procedure. In Section 3, we use extensive simulation studies to evaluate the performance of nestedIDR in comparison with the standard IDR method. In Section 4, we apply our method to an ENCODE ChIP-seq dataset (Davis *and others*, 2017; ENCODE Consortium et al., 2012) and a SEQC RNA-seq dataset (Su *and others*, 2014) and evaluate the biological relevance of the results. Section 5 is a general discussion.

## 2. Method

### 2.1 Model

Suppose L labs perform independent replications and each lab has *M_l_* (*l* = 1,…, *L*) replicates (Figure 1). For each replicate, n candidates are measured using a high-throughput experiment. A quantitative measurement is obtained for each candidate on each replicate to indicate the strength of evidence for the candidate to be a true signal of interest, e.g. gene expression level from an RNA-seq experiment or p-values of protein binding sites on DNA identified from a ChIP-seq experiment. Without loss of generality, we assume that all labs have the same number of replicates, i.e. *M*_1_ = *M*_2_ = … = *M_L_* = *M*, and denote the total number of replicates as *P* = *LM*. Let *X* be an *n* × *P* data matrix representing the measurements for the *n* candidates from the *P* replicates. Denote the measurements for the *i*^th^ candidate as a vector of length *P*, 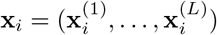, where 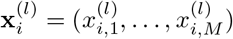 are from the *M* replicate samples from the *l*^th^ lab. We denote the distribution of the measurements on the *m^th^* replicate sample in the *l^th^* lab, 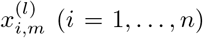, as 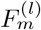, where 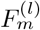 is unknown and can vary across replicates and labs. In fact, our methodology only needs the information of the ranking of the measurements on each replicate. Thus, if the actual measurements are unavailable and only the ranking information is known, 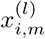 will denote the ranking of the *i^th^* candidate on the *m^th^* replicate sample and 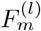 will be the empirical distribution. Without loss of generality, we assume that the candidates with a stronger measurements to be true signals receive higher ranks.

Figure 2 shows the schematic structure of our model. We assume that the data consists of both genuine signals and noise, and that genuine signals are replicable across labs but noise is not. We use *G_i_* to denote whether the *i^th^* candidate is a genuine signal (*G_i_* = 1) or noise (*G_i_* = 0), and use *π*_0_ and *π*_0_ = 1 – *π*_1_ to denote the proportions of signals and noise in the candidates, respectively. Whether or not a genuine signal is reproducible within a lab depends on the data quality of that lab and the signal strength. For example, if the sequencing depth, a key experimental factor in sequencing experiments, is sufficient in one ChIP-seq replicate but insufficient in another replicate, then some protein binding sites may be detected only in the deeper replicate and thus are irreproducible across the two replicates. Thus if the sequencing depth is sufficient for both replicates in Lab 1 but not for one of the replicates in Lab 2, then a ChIP-seq binding site can be reproducibly observed on both replicates in Lab 1, but not in Lab 2. We use 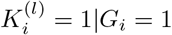 or 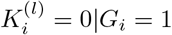 to denote whether a candidate *i* is reproducible or not in the *l^th^* lab, given that the candidate is a true signal, and use 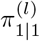 and 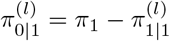 to denote the corresponding proportion in the *l^th^* lab. Because noise is not reproducible, we assume 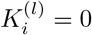 when *G_i_* = 0.

**Fig. 2.**
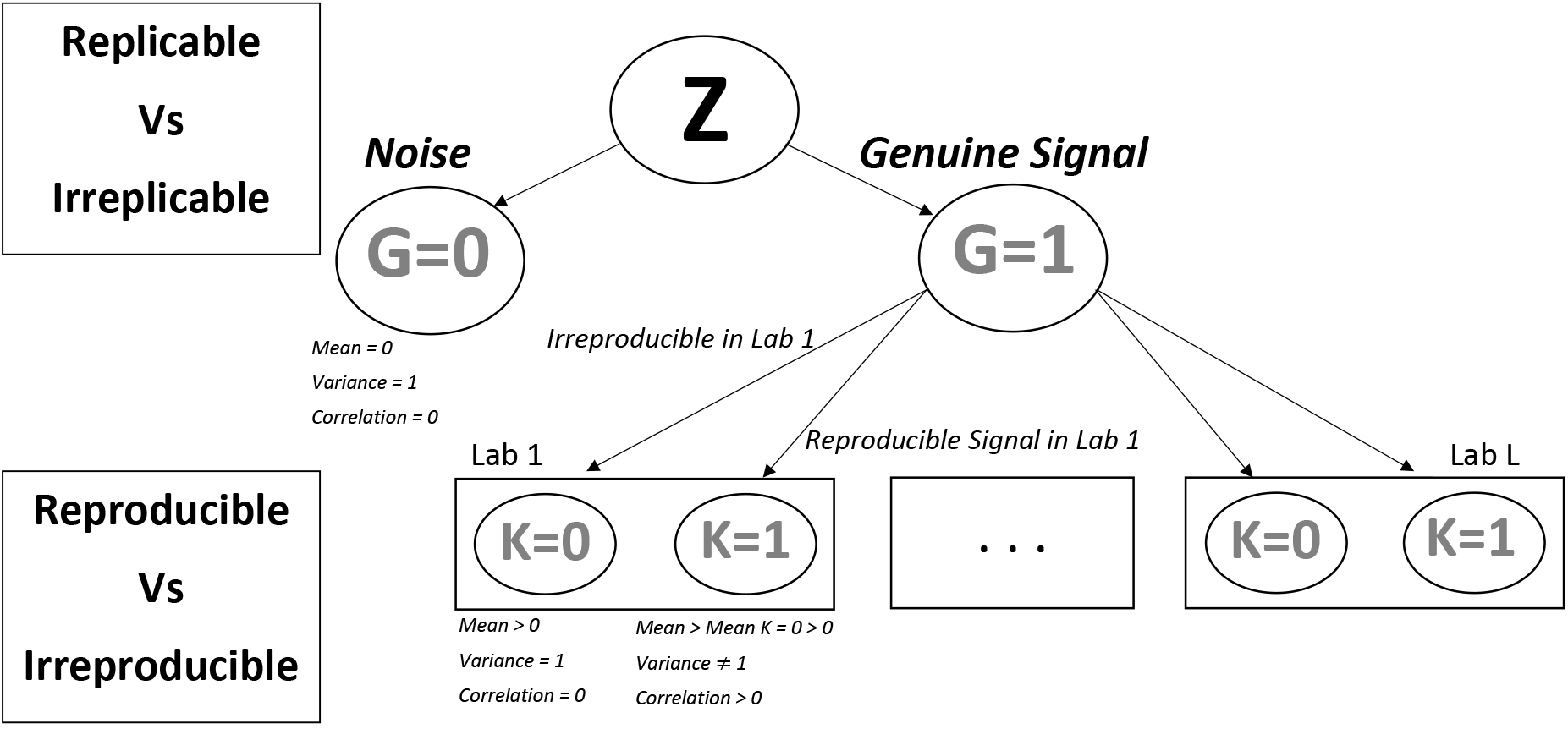
Structure of our model.

We model the data using a semiparametric copula framework as in the IDR model (Li *and others*, 2011). This framework represents the dependence structure across replicates by a simple parametric model and estimates the marginal distributions nonparametrically using ranks to permit arbitrary scales. It conveniently and robustly accommodates the situation that the measurements distribute differently across replicates or are known only by the ranking. Similar to the IDR model, we use a mixture model to model the heterogeneity of the dependence structure. However, unlike the IDR model, which assumes the measurements correlate similarly across all replicates, the dependence structure in multi-source studies is more complicated: the replicates from the same data source are more strongly correlated than those from different data sources due to source-specific effects. To model this structure, we assume that the dependence for the *i^th^* candidate is induced by a latent vector 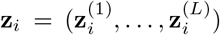, where 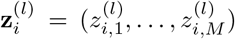 and 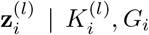, *G_i_* follows a multivariate Gaussian distribution. Because replications in different labs are done independently, 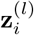 is assumed to be independent across labs. For noise (i.e. *G_i_* = 0), 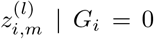 is assumed to be independent across all replicates in all labs, i.e. for all *m* ∈ {1,…, *M*} and all *l* ∈ {1,…, *L*}. For a genuine signal (i.e. *G_i_* = 1), if it is irreproducible in lab *l* (i.e. 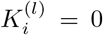), then 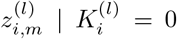, *G_i_* = 1 is assumed to be independent across replicates (*m* ∈ {1,…, *M*}) from lab *l*; whereas, if it is reproducible (i.e. 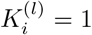), then 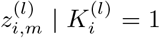, *G_i_* = 1 is assumed to have a positive correlation across replicates from this lab. Since genuine signals (i.e. *G_i_* = 1) are usually stronger than noise and reproducible signals are usually stronger than irreproducible ones, we expect that real signals have higher means than noise and reproducible signals have a higher mean than the irreproducible ones. Moreover, we assume that, for a given combination of *k* and *g*, the latent variable 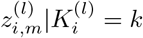, *G_i_* = *g* have the same marginal distribution across replicates from the same lab, because the underlying biology is identical and replicate samples from the same lab often share some technical commonalities. Note that this assumption on marginal distribution only applies to the latent variables *Z*, rather than the actual observations *X*. Our model still allows the marginal distribution of the actual observations to vary across replicates. Lastly, note that 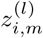 reflects the relative ordering of the data rather than the actual measurements, thus only the relative difference in means and the ratio of the variance between latent variables 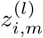 can be characterized. To ensure identifiability, we set the mean of 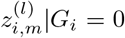 to 0 and the variances of 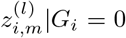 and 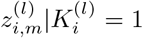, *G_i_* =0 to 1.

Taken together, we can write our model as a data generation process that consists of two steps. First, the dependence structure is generated for the measurements within Lab *l*, (*l* = 1,…, *L*), according to the status of reproducibility and replicability as

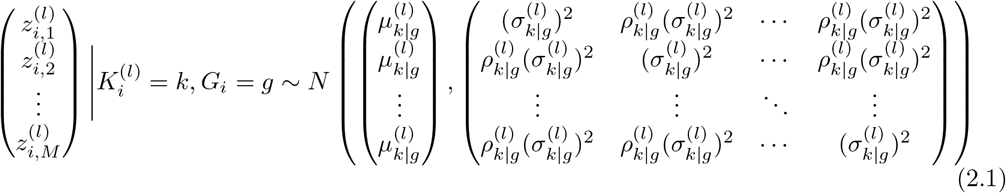

where 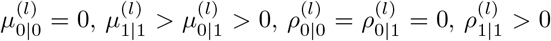, and 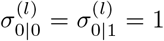. Next, the actual observation 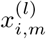 is generated by inverse probability transformation as follows,

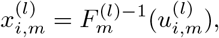

where 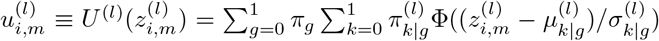 and Φ(·) is the CDF of the standard normal distribution.

The realizations, 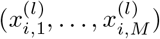, inherit the joint dependence structure from the latent vector, 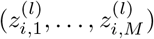. To see this more clearly, note that each 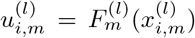 has a uniform distribution on [0,1] and the joint dependence of 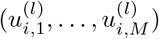 is induced from that of 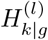, where 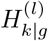 is the cumulative density function (CDF) of the distribution of 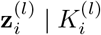, *G_i_* in (2.1). In other words, the joint CDF of 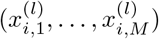 can be written as a copula that has the same dependence structure as 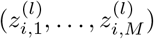 as follows:

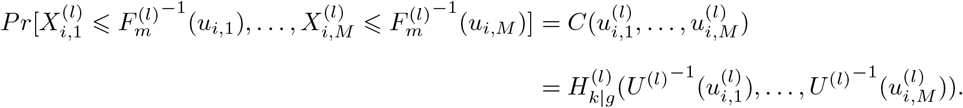

Because the dependence structure in this model is induced using a nested mixture model, we call our method nestedIDR. The IDR model can be viewed as a special case of nestedIDR with *L* = 1.

This model is parameterized by

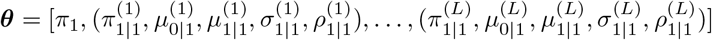

and 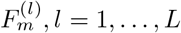. Because 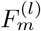 is unknown, it will be substituted by the empirical distribution. The corresponding mixture likelihood for the data is

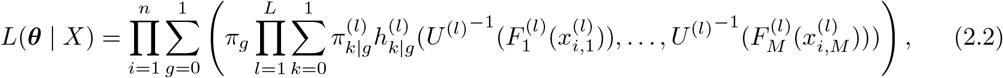

where 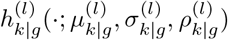 is the multivariate normal density function in (2.1) and *U*^(*l*)−1^ is the probabilistic inverse function of *U*^(*l*)^. The parameter vector ***θ*** can be estimated using an estimation procedure described in Section 2.3.

### 2.2 Measure of reproducibility and replicability

Given the parameter vector ***θ***, we can determine the replicability and the reproducibility for each candidate. To assess the replicability for a candidate *i* across different data sources, we calculate the posterior probability that the candidate belongs to the noise group (*G_i_* = 0), which is defined as

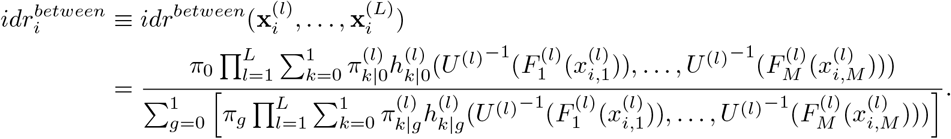

Using *idr^between^*, one can quantify the replicability across sources for each candidate. Because candidates that are highly replicated across independent sources are likely to be real signals, this quantity can be thought of as a score that judges the authentication of the signals based on the replication information aggregated from all replicates across sources.

To assess the reproducibility of a replicable candidate across replicates from the same source, we calculate the posterior probability that the candidate is irreproducible 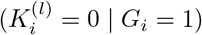 in source *l* given that it is replicable, which is defined as

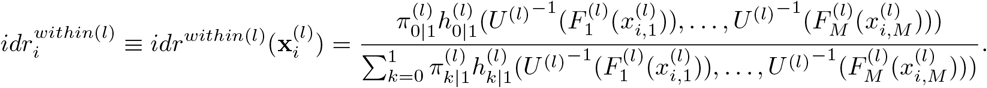

Using *idr*^*within*(*l*)^, one can identify reproducible candidates from a given source and infer the overall data reproducibility for that source.

Furthermore, we define the expected rate of irreplicable discoveries for a set of candidates selected based on *idr^between^*, analogous to that in the standard IDR framework (Li *and others*, 2011), as

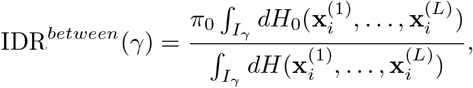

where 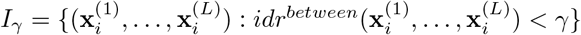, *H_g_* and *H* are the CDF of the joint density functions

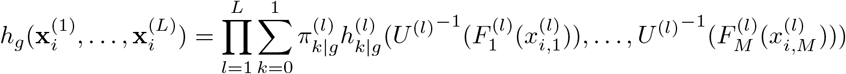

and

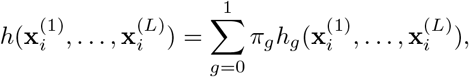

respectively. For a desired control level *α*, we denote the candidate ranked at the *i^th^* position by *idr^between^* values in an ascending order as 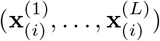 and define 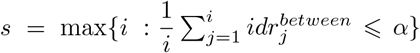. Then by selecting all 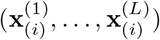 with *i* = 1,…, *s*, this procedure gives an expected rate of irreplicable discoveries no greater than *α*.

Similarly, we define the expected rate of irreproducible discoveries in a set of candidates selected based on *idr*^*within*(*l*)^ from the lab *l* as

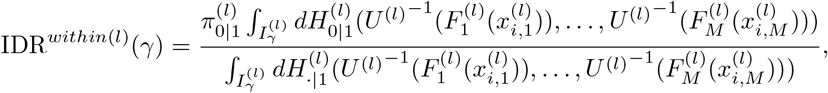

where 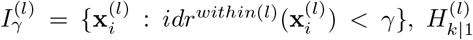, 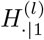 and 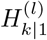 are the CDF of density functions 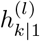 and 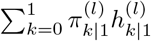, respectively. For a desired control level *α*, we denote the candidate that is ranked at the *i^th^* position by *idr*^*witthin*(*l*)^ values in an ascending order as 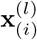 and define 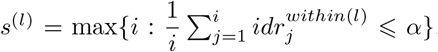. By selecting all 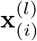 with *i* = 1,…, *s*^(*l*)^, this procedure gives an expected rate of irreproducible discoveries no greater than *α* in lab *l*.

### 2.3 Estimation

Following the approach in Li *and others* (2011), we carry out parameter estimation through a pseudo EM algorithm. Specifically, we first compute the pseudo-data 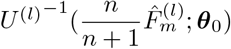 from some initialization parameters ***θ***_0_, where 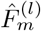 is the empirical marginal CDF and *n*/(*n* + 1) is a rescaling factor to avoid infinities. Then the algorithm iterates between two steps: (1) maximizing the likelihood (2.2) with respect to ***θ*** based on the pseudo-data using an expectation-maximization (EM) algorithm and (2) updating the pseudo-data based on the parameter estimated in (1).

To select reasonable starting values, one can first perform a preliminary exploratory data analysis to determine the likely range of values for each parameter. Then, the algorithm is run from multiple sets of starting values, and the estimates that achieve the maximum observed likelihood value are selected as the final estimates. Note that while step (1) guarantees that the likelihood (2.2) is non-decreasing for a given instance of pseudo data, the likelihood does not necessarily increase after updating the pseudo data (Bilgrau *and others*, 2016). In our experience, the observed likelihood always increases at the initial steps; however, when the algorithm is reaching convergence, it behaves similarly to the stochastic EM algorithm, i.e. the likelihood fluctuates within a narrow range (10^−2^). Thus, we stopped the algorithm whenever one of the following criteria holds: the difference between two consecutive likelihoods is small as in Li *and others* (2011), or the difference between the parameter estimates is small. The parameter estimate obtained at the highest likelihood is used as the final estimate.

## 3. Simulation Study

We conducted a simulation study to evaluate the performance of our method and compared it to existing methods. We simulated the data from the nested copula model for two labs, each of which has two replicate samples. To investigate the performance of the methods at different scenarios, we varied the signal-to-noise ratio by adjusting the proportion of noise π_0_ in the data and the level of variation (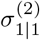 and 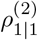) between replicate samples from a given lab. This setup produced four different simulation settings (Table 1): (S1) high signal-to-noise ratio and low variation in measurement for both labs, (S2) high signal-to-noise ratio and low variation in measurement for one lab but high for the other lab, (S3) low signal-to-noise ratio and low variation in measurement for both labs, and (S4) low signal-to-noise ratio and low variation in measurement for one lab but high for the other lab. Among them, S2 (S4, respectively) has the same signal-to-noise ratio and the same measurement variation in lab 1 as S1 (S3, respectively), but a lower measurement variation in lab 2 than S1 (S3, respectively). This simulation design allows us to compare the effect due to the signal-to-noise ratio of specimen distributed to all labs and the effect due to reproducibility issues associated with an individual lab. For each simulation setting, we generated 200 datasets each with *n* =5000 candidates.

**Table 1.** True parameters for four simulation settings.

| Scenario | $\pi_0$ | $\pi_{0\vert 1}^{(1)}$ | $\pi_{0\vert 1}^{(2)}$ | $\mu_{0\vert 1}^{(1)}$ | $\mu_{0\vert 1}^{(2)}$ | $\mu_{1\vert 1}^{(1)}$ | $\mu_{1\vert 1}^{(2)}$ | $\sigma_{1\vert 1}^{(1)}$ | $\sigma_{1\vert 1}^{(2)}$ | $\rho_{1\vert 1}^{(1)}$ | $\rho_{1\vert 1}^{(2)}$ |
| --- | --- | --- | --- | --- | --- | --- | --- | --- | --- | --- | --- |
| S1 | 0.3 | 0.2 | 0.2 | 1 | 1 | 3 | 3 | 1 | 1.0 | 0.9 | 0.9 |
| S2 | 0.3 | 0.2 | 0.5 | 1 | 1 | 3 | 3 | 1 | 1.4 | 0.9 | 0.7 |
| S3 | 0.5 | 0.2 | 0.2 | 1 | 1 | 2 | 2 | 1 | 1.0 | 0.9 | 0.9 |
| S4 | 0.5 | 0.2 | 0.5 | 1 | 1 | 2 | 2 | 1 | 1.4 | 0.9 | 0.7 |

We compared our method with the standard IDR model by running the standard IDR model in two ways: (1) using all four replicates from both labs (referred to as *idr^a^*) and (2) using only the replicates from a single lab (referred to as *idr*^*l*_1_^ for lab 1 and *idr*^*l*_2_^ for lab 2). Since the former measures the concordance among replicates across labs, it functions as a measure of replicability, i.e. a counterpart of *idr^between^* in nestedIDR. The latter only involves replicates from a single lab, thus reflecting the reproducibility within a lab and functioning as a baseline comparison with *idr^between^*. These comparisons allow us to assess and quantify the importance of modeling heterogeneity across different sources.

### 3.1 Results

#### 3.1.1 Assessment of replicability

Replicability is indicative of the likelihood of a signal being real. We thus evaluated the performance of each method in assessing replicability by its sensitivity and specificity for distinguishing true signals from noise at a series of thresholds (see Table 2 and Figure 3). In all settings, the methods using replicates from both labs (nestedIDR and *idr^a^*) show higher area-under-the-curve (AUC) than those only using the replicates from a single lab. When the data quality is different between labs (S2 and S4), the performance of the latter heavily depends on the data quality of the specific lab (AUC (*idr*^*l*_1_^ vs *idr*^*l*_2_^): 0.965 vs 0.858 for S2, and 0.918 vs 0.839 for S4, respectively), whereas the methods integrating data from all labs still perform robustly. This shows that data integration across different sources can improve the reliability of identification.

**Fig. 3.**
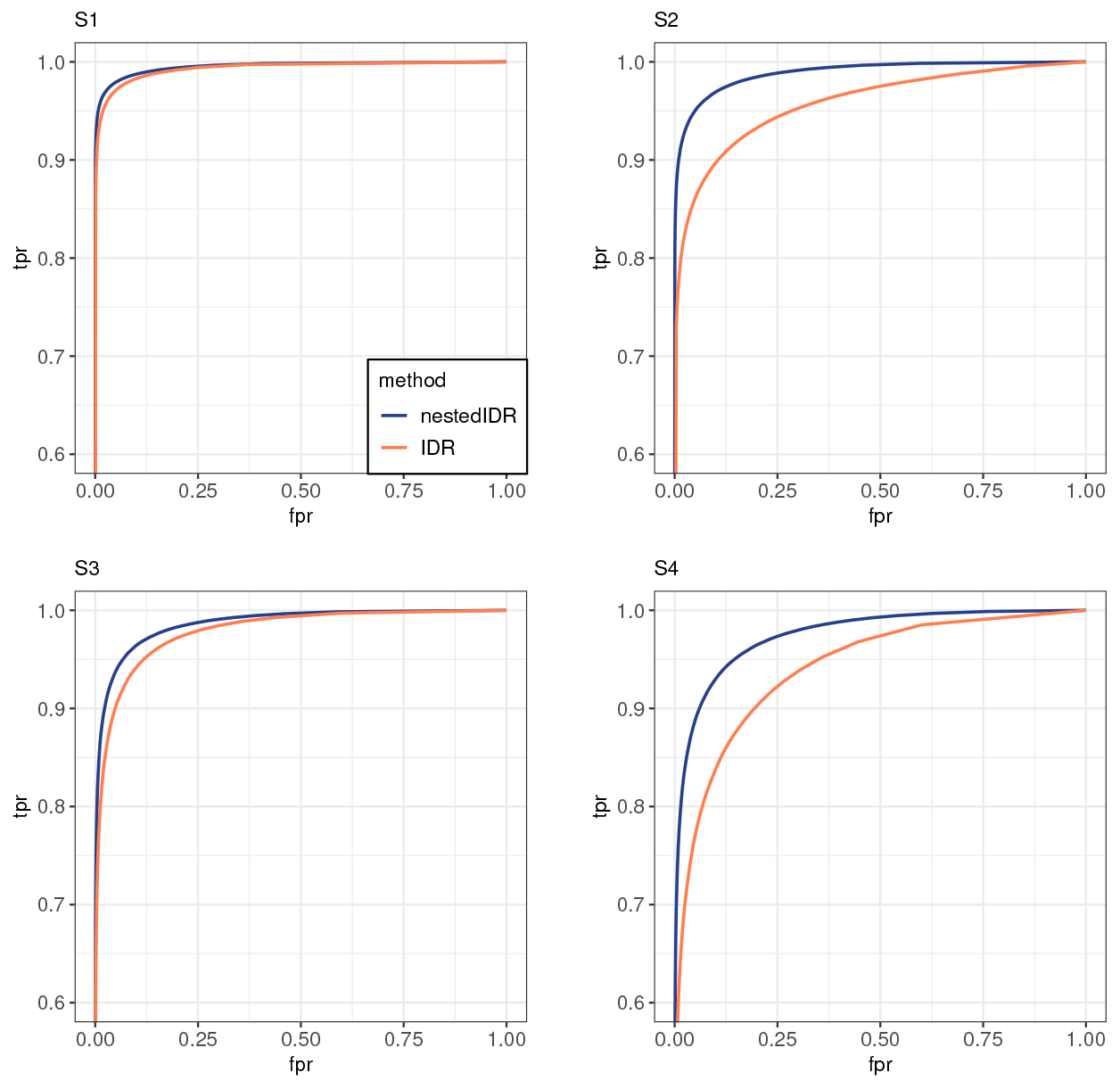
Comparison of ROC curves at different simulation settings. Blue solid: Nested IDR(*idr^between^*); Orange solid: IDR (*idr^a^*). Each ROC curve is the average over 200 simulation datasets.

**Table 2.**
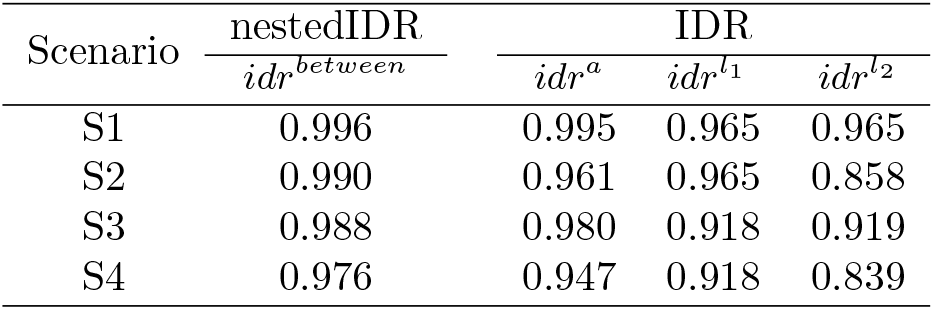
Comparison of AUC values in the four simulation settings.

| Scenario | nestedIDR | IDR |  |  |
| --- | --- | --- | --- | --- |
| | $idr^{between}$ | $idr^a$ | $idr^{l_1}$ | $idr^{l_2}$ |
| S1 | 0.996 | 0.995 | 0.965 | 0.965 |
| S2 | 0.990 | 0.961 | 0.965 | 0.858 |
| S3 | 0.988 | 0.980 | 0.918 | 0.919 |
| S4 | 0.976 | 0.947 | 0.918 | 0.839 |

When data quality is similar between two labs (S1 and S3), our method and *idr^a^* perform quite similarly (AUC (nestedIDR vs. *idr^a^*): 0.995 vs. 0.993 for S1 and 0.986 vs. 0.978 for S3, respectively). However, when data quality is different between labs (S2 and S4), our method outperforms *idr^a^* (AUC (nestedIDR vs. *idr^a^*): 0.989 vs. 0.957 for S2 and 0.975 vs. 0.940 for S4, respectively). This is because our method accounts for heterogeneity in degrees of reproducibility of data from different labs, whereas the standard IDR model is source-agnostic.

#### 3.1.2 Comparison of classification accuracy

Because replicable signals are identified as the candidates that exceed a prespecified replicability cutoff, we next evaluate the classification accuracy for the replicable signals identified at a given cutoff. Here, for nestedIDR, we classified a candidate as replicable if *IDR^between^* ≼ 0.05 and irreplicable otherwise for each sample. For the methods based on the standard IDR (i.e. *idr^a^, idr*^*l*_1_^ and *idr*^*l*_2_^), we classified candidates similarly according to their corresponding IDR values. We then used Adjusted Rand Index (ARI) (Hubert, 1985), a commonly used index for measuring the agreement between clusterings, to compare the classification from each method with the true classification. Given two clusterings, ARI has the expected value of 0 if the clusterings are independent, and it reaches the maximum value of 1 if they are identical.

Figure 4 shows the distribution of ARI for each simulation setting. For all settings, our method has the highest classification accuracy among all methods. In addition, its ARI also has a much smaller variation across simulation datasets than *idr^a^*, especially when the replicates are heterogeneous, for example, S2 (SD of ARI: 0.012 for nestedIDR and 0.150 for *idr^a^*) and S4 (SD of ARI: 0.012 for nestedIDR and 0.136 for *idr^a^*). The observed high variation of classification accuracy in *idr^a^* is likely related to the homogeneous treatment of all replicates in the standard IDR method: once the homogeneous assumption is violated, its parameter estimation becomes less stable, undermining its quality and stability of classification. On the other hand, by accounting for the heterogeneity across labs, nestedIDR effectively maintains the quality and stability of its classification in these scenarios.

**Fig. 4.**
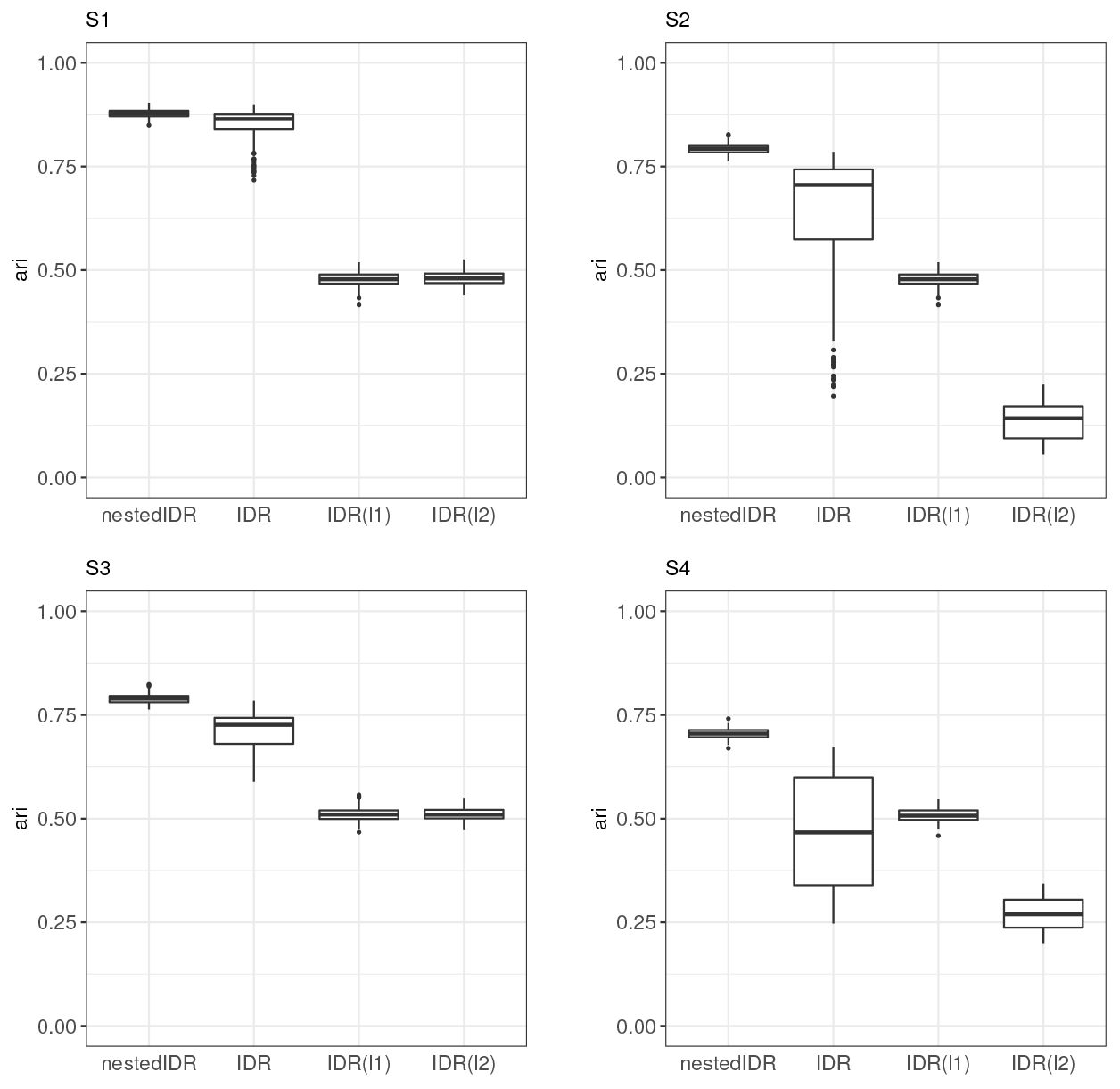
Comparison of classification accuracy measured by ARI in the simulations S1-S4.

#### 3.1.3 NestedIDR can distinguish between different causes of discordance

Next we assessed the utility of our method as a tool for diagnosing data quality issues in multi-lab studies. In particular, we evaluated whether our method could properly detect the reduction of concordance and illuminate the cause of said reduction when the concordance of findings across replicates is diminished. Here we focused on two common causes that can reduce the concordance of findings in multi-lab studies: (1) low signal-to-noise ratios in the specimen distributed to all labs and (2) high variation in the measurement procedure of a lab. The former makes replication harder both within and across labs, thereby reducing both reproducibility and replicability. However, if the latter occurs in one lab, this has a larger impact on the reproducibility of the affected lab and a smaller impact on the overall replicability.

To illustrate scenario (1), we consider the change from S1 to S3 and from S2 to S4. They represent the cases that the signal-to-noise ratio is reduced but the level of variation in the measurement procedure is intact for each lab, where the two labs have similar (or different, respectively) variation in S1-S3 (S2-S4, respectively). To illustrate scenario (2), we consider the change from S1 to S2 (or from S3 to S4, respectively). They represent the cases that the variation of lab 2 increases but the signal-to-noise ratio maintains the same high (or low, respectively) level. We compare the performance of nestedIDR and the standard IDR methods. To visualize the change of reproducibility or replicability at different cutoffs, we plotted the number of candidates selected by nestedIDR and the standard IDR method at a series of cutoffs (Figure 5). Given a replicability (or reproducibility) cutoff, a larger number of selected candidates indicates higher replicability (or reproducibility) of the data.

**Fig. 5.**
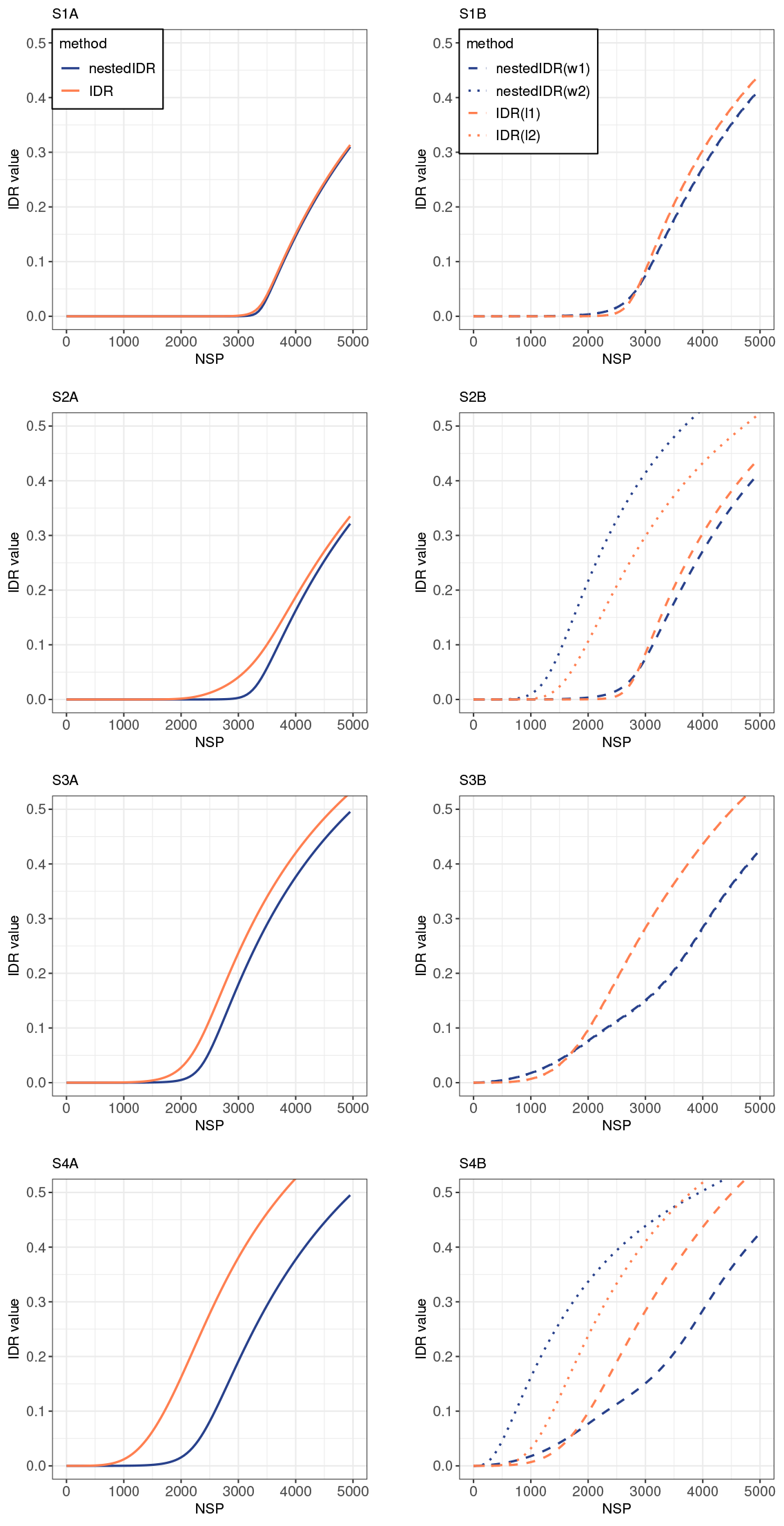
The number of selected signals at different reproducibility and replicability cutoffs in the four simulation settings. The left column shows the number of *replicable* signals selected using nestedIDR (*idr^between^*, blue solid) and IDR (*idr^a^*, orange solid) with data from both labs. The right column shows the number of *reproducible* signals selected using nestedIDR and IDR with the data from a single lab. nestedIDR(w1): lab 1 results from *idr*^*within*(1)^ (blue dashed); nestedIDR(w2): lab 2 results from *idr*^*within*(2)^ (blue dotted); IDR(l1): lab 1 results from *idr*^*l*_1_^ (orange dashed); and IDR(l2): lab 2 results from *idr*^*l*_2_^ (orange dotted).

When the signal-to-noise ratio is lowered and the variation of measurement procedures remains intact, both nestedIDR (*idr^between^, idr*^*within*(*l*_1_)^ and *idr*^*within*(*l*^2^)^) and the standard IDR methods (*idr^a^, idr*^*l*_1_^ and *idr*^*l*_2_^) select substantially fewer signals with a reduction of 31.0-41.8% at the cutoff of 0.05 (Figure 5 S1 vs S3, or Figure 5 S2 vs S4), correctly reflecting that the concordance is compromised beyond individual labs. When the signal-to-noise ratio is intact but the variation of measurement procedures from lab 2 increases (Figure 5 S1 vs S2 and Figure 5 S3 vs S4), the number of reproducible signals identified by *idr*^*within*(*l*_2_)^ is substantially decreased (53.4% for S1 vs S2 and 68.4% for S3 vs S4), whereas the number of reproducible signals identified by *idr*^*within*(*l*_1_)^ from lab 1 is unchanged and the numbers of replicable signals identified by *idr^between^* is only slightly reduced (3.14% and 5.68%, respectively). This pattern signals that the lowered concordance is due to low reproducibility in lab 2, rather than the compromised signal-to-noise ratio. In contrast, while the standard IDR-based method correctly reports lowered reproducibility in lab 2 (*idr*^*l*_2_^), the number of replicable signals (identified by *idr^a^*) is also greatly reduced (12.7% for S1 vs S2 and 33.3% for S3 vs S4), showing a pattern indifferent from the scenario of lowered signal-to-noise ratio. Together, this demonstrates the effectiveness of nestedIDR for differentiating the causes of reduced concordance and diagnosing data quality issues.

## 4. Applications to real data

### 4.1 Application to the ENCODE ChlP-seq datasets

In our first analysis, we obtained ChIP-seq data from the ENCODE project (ENCODE Consortium et al., 2012). According to ENCODE experimental design, for each binding protein, independent ChIP-seq experiments were performed in two or more labs, with at least two biological replicates from each lab. Here we assessed the performance of nestedIDR using the ChIP-seq experiments for the transcription factor CTCF on the K562 cell line. The data were generated by two labs, University of Texas at Austin (UTA) and University of Washington (UW), with two replicates contributed from each lab (UTA replicate 1: 4,861,426 reads, UTA replicate 2: 4,940,148 reads, UW replicate 1: 12,853,025 reads and UW replicate 2: 5,676,639 reads). We refer to UTA as lab 1 (l1) and UW as lab 2 (l2) in our analyses. We downloaded the BAM files directly from the ENCODE data portal (Davis *and others*, 2017; Landt *and others*, 2012) and called ChIP-seq peaks using MACS2 (Feng *and others*, 2012; Zhang *and others*, 2008) for each replicate sample. Due to variation across samples, the locations of ChIP-seq peaks identified for the same binding event may be slightly different across replicates. We therefore merged the ChIP-seq peaks that overlap across replicates using the bedtools (Quinlan and Hall, 2010) command `“intercept”`. For each merged region, we used -log10(p-value) calculated by MACS2 at the highest peaks within a merged region to represent the signal for the region. Then the reproducibility and replicability of the peaks across the replicates were evaluated using nestedIDR and the standard IDR methods based on these signals.

#### 4.1.1 NestedIDR provides more consistent results with motif binding scores

To validate the biological relevance of our results, we evaluated the motif occurrence at the identified ChIP-seq peaks. A motif refers to a short DNA sequence that is preferentially bound by a binding protein. While true binding sites do not necessarily contain motifs, they generally have a higher frequency of motif occurrence than other regions. Here we obtained the FIMO score (Grant *and others*, 2011) for the binding sites identified in the ChIP-seq experiments, which measures the likelihood of motif occurrence for each binding site, using the MEME suite (Bailey *and others*, 2009). Because signals in ChIP-seq experiments reflect the experimentally measured likelihood of protein binding, the replicability of the ChIP-seq peaks is expected to be positively correlated to their FIMO scores. We therefore evaluated the biological relevance of the replicability assessment by computing the Spearman correlation between the FIMO score and the replicability score assigned by each method.

We observed that the results from nestedIDR have the higher Spearman correlation with the FIMO score (*idr^between^*: 0.440) than all the standard IDR methods (*idr^a^*: 0.425, *idr*^*l*_1_^: 0.413 and *idr*^*l*_2_^: 0.388). This demonstrates the effectiveness of nestedIDR in identifying biologically relevant signals.

#### 4.1.2 NestedIDR can rescue signals that are irreproducible in a single lab but are replicated by the other lab

An important benefit of multi-source analyses is the possibility to recalibrate the confidence of identifications made in single sources using the replication information across sources. For example, if a putative binding site that shows moderate reproducibility within a lab is replicated by another lab, the confidence of identification will be greatly boosted. To evaluate the effectiveness of using replication information across labs, we compared the sets of peaks that are identified as replicable by nestedIDR and the standard IDR. Because both methods are mixture models, these sets cab be defined using the posterior probabilities (*idr^between^* and *idr^a^*) of belonging to the replicable group. Here we defined the nestedIDR-specific set as the peaks satisfying *idr^between^* < 0.3 and *idr^a^* > 0.6 and the IDR-specific set as the peaks satisfying *idr^between^* > 0.6 and *idr^a^* < 0.3, respectively. We left out the peaks with the posterior probabilities of 0.3-0.6 to ensure the confidence of the specific identifications. There are 710 peaks specifically identified by nestedIDR and 773 specifically identified by the standard IDR. As shown in Figure 6A, though the range of the FIMO scores is similar between these two sets, the proportion of peaks with highly significant FIMO scores is notably higher in the nestedIDR-specific set than the IDR-specific set. For example, 39.52% nestedIDR-specific peaks have FIMO scores < 10^−6^ in comparison with 30.70% for IDR-specific peaks.

**Fig. 6.**
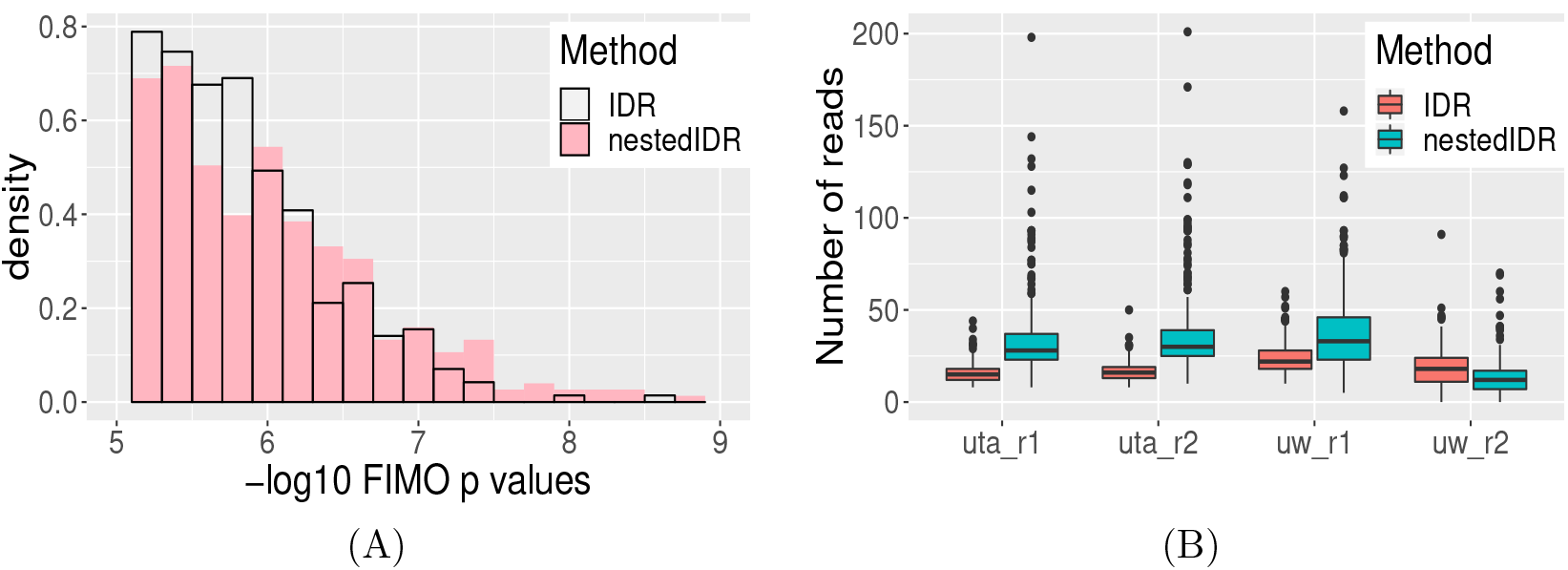
Comparison of the ChIP-seq peaks that are specifically identified by nestedIDR (*idr^between^*) and IDR (*idr^a^*). (A) The distribution of FIMO -log10(p-value) for peaks specifically identified by each method. (B) The distribution of read counts for the peaks specifically identified by each method in the four replicates.

Next, we examined the characteristics of the peaks in each set. We plotted the distributions of read counts for the two peak sets (Figure 6B). Interestingly, the IDR-specific peaks have similar read counts across replicates, whereas the nestedIDR-specific peaks have substantially higher read counts in the three replicates with strong signals (UTA-r1, UTA-r2 and UW-r1) but lower read counts in the replicate with weak signals (UW-r2) than IDR-specific peaks. To understand the characteristics of peaks in each set, we plotted five exemplary peaks in each set on the IGV genome browser (Robinson, 2011). As shown in Figure 7a, the IDR-specific peaks have relatively consistent signals across all replicates from both labs. However, their average signal strengths tend to be lower than those in the nestedIDR-specific set (Figure 7B). In contrast, the nestedIDR-specific peaks tend to be highly variable between replicate samples within one lab but are replicated with strong signals in the other lab (Figure 7B). Indeed, these peaks are deemed as highly irreproducible in the lab that shows high variation by the standard IDR based on the single-lab data. However, despite being replicated in the other lab, they are deemed as highly irreplicable across labs by *idr^a^* (Table 3). Thus they would be excluded from downstream analyses by the standard IDR. However, they receive highly favorable scores (*idr^between^*) from nestedIDR, and thus are deemed replicable by nestedIDR. These observations on the exemplary peaks explain the observed difference in the distribution of read counts between the two specifically identif ied sets (Figure 6B). Together, this shows that nestedIDR is able to rescue peaks that are weak in the low-quality replicate (UW-r2) but are replicated with strong signals in other replicates (UTA and UW-r1) by leveraging replication information across labs.

**Fig. 7.**
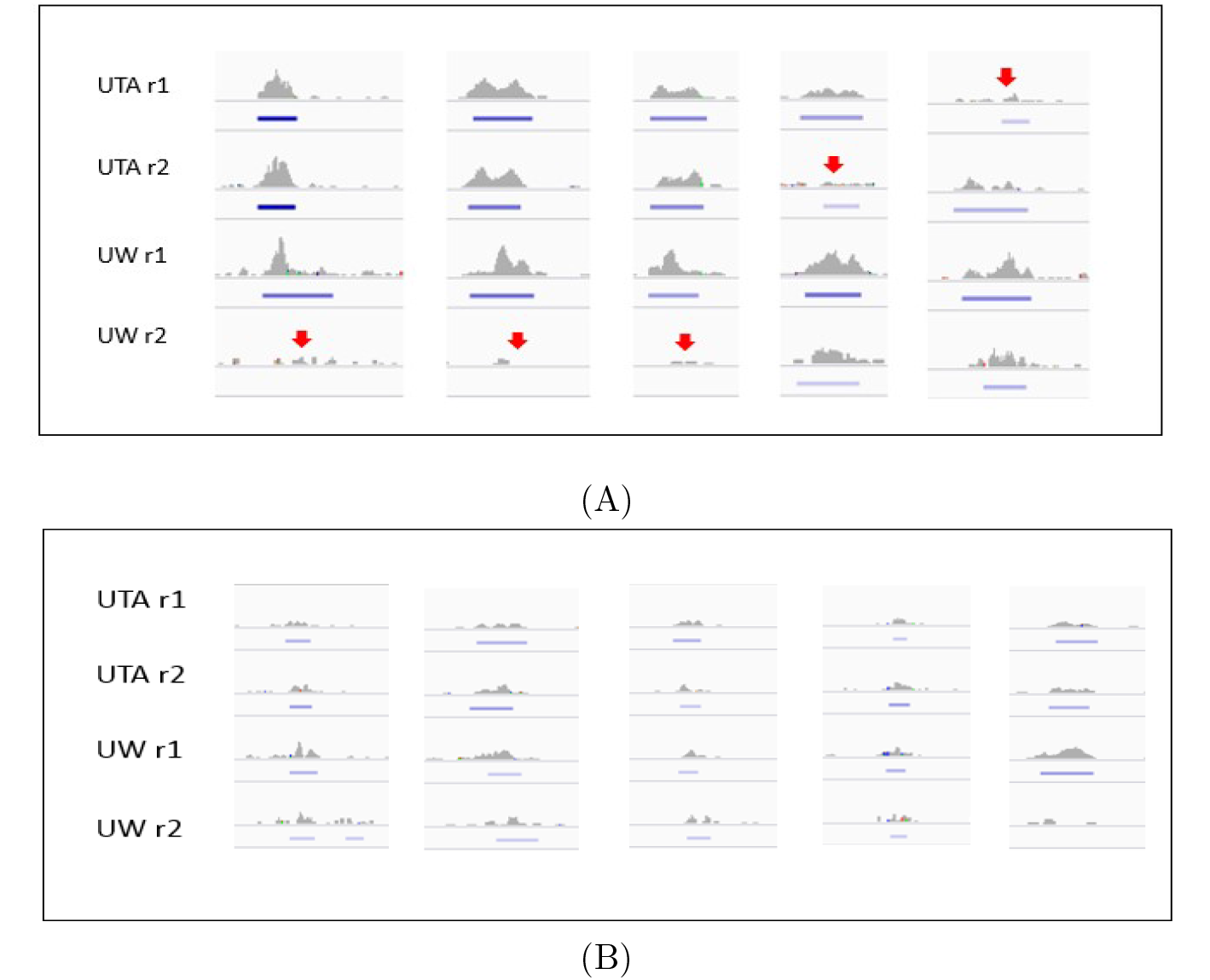
IGV Genome browser shots of example peaks (A) that are specifically identified by nestedIDR and (B) that are specifically identified by IDR (*idr^a^*). The ChIP-seq data were obtained from two labs, UTA and UW. The Y axis shows the number of mapped reads, ranged 0-40. The blue bar under each peak shows MACS2 peak calling results, where darker color indicates more significant MACS2 p-values and no bar indicates that a peak is not called by MACS2. The red arrow marks the weakest peaks among the 4 replicates.

**Table 3.**
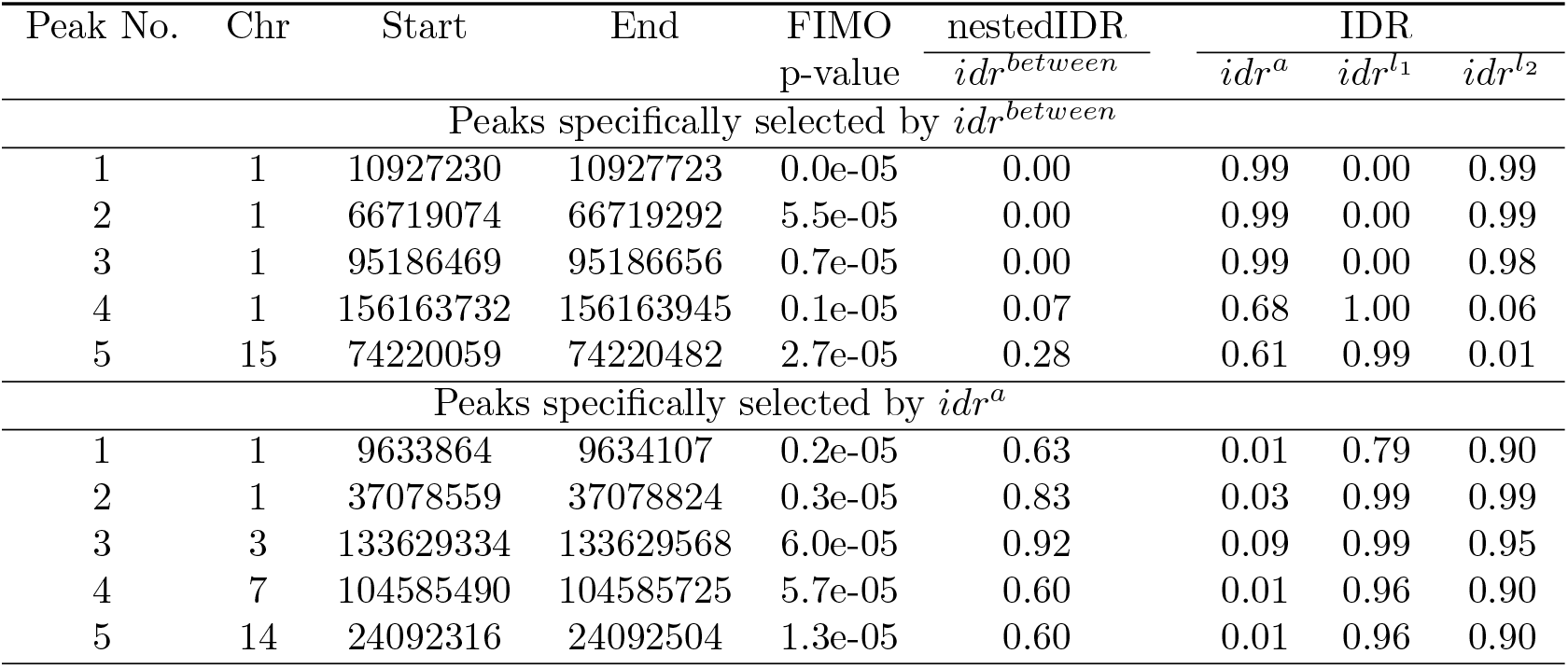
Information of the example ChIP-seq peaks in Figure 7. Lab 1 is UTA and Lab 2 is UW.

The difference between the selections of nestedIDR and the standard IDR is likely attributed to the difference in their assumptions. The standard IDR model assumes that all replicates come from a homogeneous source, so it strongly favors peaks that are consistent across all replicates. This property grants the standard IDR advantages when data are truly homogeneous, allowing it to identify moderate but consistent signals. However, when integrating data from different sources, this property hampers the identification of signals which are not consistent across all data sources. On the other hand, nestedIDR accommodates heterogeneity from multiple sources, allowing it to distinguish real signals that are irreproducible due to measurement variation in one source from false signals that are not replicable in multiple sources.

### 4.2 Application on the SEQC RNA-seq data

Next we applied nestedIDR to the RNA-seq data from the Sequencing Quality Control (SEQC) project (Su *and others*, 2014). SEQC compared the performance of three RNA-seq platforms using benchmark datasets for Universal Human Reference RNA sample and Human Brain RNA sample (see Figure 8). Each RNA sample was sequenced using three platforms, namely, Illumina HiSeq 2000, Life Technologies and Roche 454. Illumina HiSeq 2000 was used to generate data at the Mayo Clinic (MAY), BGI, Cornell, City of Hope, Novartis and the Australian Genome Research Facility. Life Technologies was used to generate data at Penn State (PSU), Northwestern (NWU), SeqWright Inc. (SQW), and Liverpool. Lastly, Roche 454 was used to generate data at the Medical Genomes Project, New York University Medical Center (NYU), and SeqWright Inc. The consortium also measured the gene expression levels for a subset of 1,129 genes using the qRT-PCR experiment, which often is deemed as a gold standard to evaluate the accuracy of RNA measurements.

**Fig. 8.**
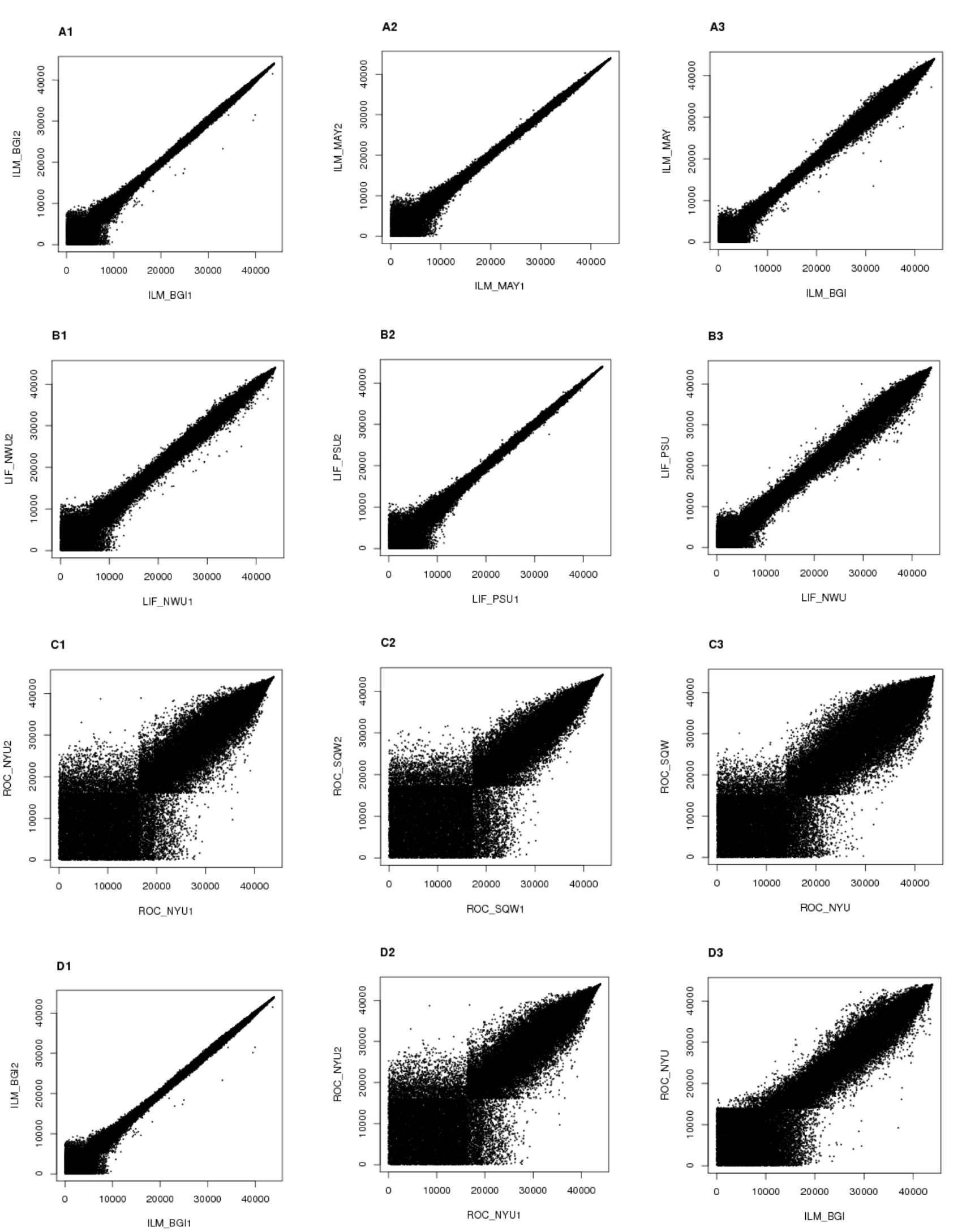
SEQC RNA-seq data. Plotted are the rank scatterplots of the gene expression level (read counts) measured using different technologies (Illumina (ILM), Life technologies (LIF) and Roche (ROC)) in different labs. Rows A-C: within-platform analysis; Row D: across-platforms analysis. Specifically, A: ILM (BGI) vs ILM (MAY); B: LIF (NWU) vs LIF (PSU); C: ROC (NYU) vs ROC (SQW); D: ILM (BGI) vs ROC (NYU). Columns 1-2: two replicates from each participating labs. Column 3: the average expression level of the two labs in the first two columns. The average expression level is calculated as the average between the two replicates in each lab.

Here we assessed the replicability and reproducibility of the gene expression levels using the Universal Human Reference RNA sample. Our analyses focus on evaluating the between-lab replicability and within-lab reproducibility for each platform. In particular, for each platform, we selected two labs (Table 4 rows 1-3), and within each lab, we used two replicate samples for data from each platform. Figure 8A-C shows the rank scatter plots of the read counts between replicate samples from the same lab (Figure 8A1-2, B1-2 and C1-2) and from different labs (Figure 8A3, B3 and C3). As shown, Illumnia and Life technologies consistently show much higher reproducibility between replicate samples than Roche. We applied nestedIDR and the standard IDR to the read counts generated from each RNA-seq platform. In addition, to evaluate the robustness of our method in presence of high heterogeneity, we also performed an across-platform analysis (Figure 8D) by comparing data generated from Illumina HiSeq 2000 at BGI with that from Roche 454 at NYU (Table 4 row 4). Similar to the within-platform analysis, we used two replicate samples within each lab. Because both lab and platform are different between the two replicate samples in the across-platform analysis, the replicates are more heterogeneous than those in the within-platform analysis.

**Table 4.** Spearman rank correlation between replicability metrics and the PCR measurements.

| Platform | Labs | nestedIDR | IDR |  |  |
| --- | --- | --- | --- | --- | --- |
| | | $idr^{between}$ | $idr^a$ | $idr^{l_1}$ | $idr^{l_2}$ |
| Illumina | BGI | 0.8593 | 0.8189 | 0.8164 | 0.8148 |
| Illumina | MAY |  |  |  |  |
| Life Technologies | PSU | 0.8903 | 0.8470 | 0.8768 | 0.8801 |
| Life Technologies | NWU |  |  |  |  |
| Roche 454 | NYU | 0.8717 | 0.8378 | 0.8571 | 0.8221 |
| Roche 454 | SQW |  |  |  |  |
| Illumina | BGI | 0.8854 | 0.8006 | 0.8164 | 0.8571 |
| Roche 454 | NYU |  |  |  |  |

#### 4.2.1 NestedIDR shows a high correlation with expression levels measured by PCR

To evaluate the biological relevance of our assessments, we compared the results from each method with the PCR measured expression value. Although PCR measurements are only available for a small subset of the genes considered in the RNA-seq datasets (1,129 out of more than 40,000 transcripts), they nevertheless provide some insights. Because an authentic signal is expected to be more replicable and stronger than noise, the ranking provided by a good replicability metric is expected to be reasonably concordant with the ranking of the true signal strength. Based on this intuition, we computed the Spearman rank correlation between the estimated metrics and PCR measurements.

As shown in Table 4, nestedIDR has the highest Spearman rank correlation with the PCR measurements in all cases. We observed that *idr^a^* sometimes performs worse than both single-lab results (*idr*^*l*_1_^ and *idr*^*l*_2_^) (Table 4 rows 2 and 4), especially when different technologies are used in the two labs (Table 4 row 4). This is likely because the homogeneous assumption in the standard IDR method (*idr^a^*) is violated when there is a high level of between-lab heterogeneity. In this situation, the integration across labs using the standard IDR can lead to results that are even worse than those obtained by only using the data from a single lab. In contrast, nestedIDR consistently outperforms the single-lab results in all cases, even when the data come from different platforms (Table 4 row 4), showing the robustness and effectiveness of nestedIDR in integrating data from heterogeneous sources.

## 5. Conclusions

In this paper, we developed nestedIDR, a nested copula mixture model, for assessing the reproducibility and replicability of high-throughput genomic data from multiple sources. By modeling the dependence between replicate samples according to their origins in a nested model, it captures the dependence structure induced by the nested experimental design in many consortium studies or studies with independent replications. This modeling approach effectively accounts for the heterogeneity across data sources and simultaneously estimates the reproducibility and replicability within and between data sources.

As illustrated in our ChIP-seq and RNA-seq data analyses, the replicability score provided by nestedIDR is more consistent with the true signal strength than the standard IDR score. It also shows a better tolerance for signals that are less reproducible within labs but reasonably replicable across labs. As a result, it is able to rescue the biologically relevant signals that are not reproducible within a data source but are replicated in other independent data sources. Finally, the proposed method is suitable not only for the genomic studies analysed here but also for other scenarios that high-throughput screenings are conducted, such as identifying signals with active brain activities in fMRI images.

## 6. Software

The R code, a sample input data set and complete documentation are available on GitHub (https://github.com/Yafei611/nestedIDR).

## 7. Supplementary Materials

Supplementary materials are available online.

