## Supplementery note & figures for "A statistical framework for measuring reproducibility and replicability of high-throughput experiments from multiple sources"

### Supplementary material to: A statistical framework for measuring reproducibility and replicability of heterogeneous omics data from multiple sources

MONIA RANALLI[†1], YAFEI LYU[†2], HILLARY KOCH[1], QUNHUA LI[*1,2]

[1]*Department of Statistics, Pennsylvania State University, University Park, Pennsylvania, United States of America*

[2]*Bioinformatics and Genomics program, the Huck Institute of the Life Science, Pennsylvania State University, University Park, Pennsylvania, United States of America*

#### Appendix - EM algorithm

1. Perform a preliminary exploratory analysis to determine the possible range of values for each parameter ($\boldsymbol{\theta}_0$).

2. Run the following algorithm from multiple starting values, and select the estimates that achieve the maximum observed likelihood value as the final estimates.

3. Compute the pseudo-data ${U^{(l)}}^{-1}(\frac{n}{n+1}\hat{F}_m^{(l)}; \boldsymbol{\theta}_0)$ from some initialization parameters $\boldsymbol{\theta}_0$, where $\hat{F}_m^{(l)}$ is the empirical marginal CDF and $n/(n+1)$ is a rescaling factor to avoid infinities.

[*]To whom correspondence should be addressed. [†]Co-first authors.

4. Estimation. The mixture likelihood for data is given by

$$L(\boldsymbol{\theta}) = \prod_{i=1}^{n}\sum_{g=0}^{1}\left(\pi_g \prod_{l=1}^{L}\sum_{k=0}^{1}\pi_{k|g}^{(l)} h_{k|g}^{(l)}({U^{(l)}}^{-1}(F_1^{(l)}(x_{i,1}^{(l)})),\ldots,{U^{(l)}}^{-1}(F_M^{(l)}(x_{i,M}^{(l)})))\right),$$

where $h_{k|g}^{(l)}$ is the multivariate normal density function with the parameters $\mu_{k|g}^{(l)}$, $\sigma_{k|g}^{(l)}$, and $\rho_{k|g}^{(l)}$. The parameter vector $\boldsymbol{\theta}$ can be estimated using a pseudo likelihood estimation. Furthermore, we assume that $\mu_{0|0}^{(l)} = 0$, $\mu_{1|1}^{(l)} > \mu_{0|1}^{(l)} > 0$, $\rho_{0|0}^{(l)} = \rho_{0|1}^{(l)} = 0$, $\rho_{1|1}^{(l)} > 0$, and $\sigma_{0|0}^{(l)} = \sigma_{0|1}^{(l)} = 1$.

Thus the complete log-likelihood is given by

$$\begin{aligned}\ell_c(\boldsymbol{\theta}) = &\sum_{i=1}^{n}\sum_{g=0}^{1} w_{ig}\log\pi_g + \sum_{i=1}^{n}\sum_{g=0}^{1}\sum_{l=1}^{L}\sum_{k=0}^{1} w_{ig}w_{ik|g}^{(l)}\log\pi_{k|g}^{(l)} + \\ &+\sum_{i=1}^{n}\sum_{g=0}^{1}\sum_{l=1}^{L}\sum_{k=0}^{1} w_{ig}w_{ik|g}^{(l)}\log h_{k|g}^{(l)}({U^{(l)}}^{-1}(F_1^{(l)}(x_{i,1}^{(l)})),\ldots,{U^{(l)}}^{-1}(F_M^{(l)}(x_{i,M}^{(l)})))\end{aligned} \tag{0.2}$$

It follows that the EM algorithm is constructed as follows.

- E-step

$$w_{ig} = \frac{\pi_g \prod_{l=1}^{L}\sum_{k=0}^{1}\pi_{k|g}^{(l)}h_{k|g}^{(l)}({U^{(l)}}^{-1}(F_1^{(l)}(x_{i,1}^{(l)})),\ldots,{U^{(l)}}^{-1}(F_M^{(l)}(x_{i,M}^{(l)}))))}{\sum_{g=0}^{1}\pi_g \prod_{l=1}^{L}\sum_{k=0}^{1}\pi_{k|g}^{(l)}h_{k|g}^{(l)}({U^{(l)}}^{-1}(F_1^{(l)}(x_{i,1}^{(l)})),\ldots,{U^{(l)}}^{-1}(F_M^{(l)}(x_{i,M}^{(l)}))))}$$

$$w_{ik|g}^{(l)} = \frac{\pi_{k|g}^{(l)}h_{k|g}^{(l)}({U^{(l)}}^{-1}(F_1^{(l)}(x_{i,1}^{(l)})),\ldots,{U^{(l)}}^{-1}(F_M^{(l)}(x_{i,M}^{(l)}))))}{\sum_{k=0}^{1}\pi_{k|g}^{(l)}h_{k|g}^{(l)}({U^{(l)}}^{-1}(F_1^{(l)}(x_{i,1}^{(l)})),\ldots,{U^{(l)}}^{-1}(F_M^{(l)}(x_{i,M}^{(l)}))))}$$

- M-step

$$\hat{\pi}_g = \frac{\sum_{i=1}^{n} w_{ig}}{n}$$

$$\hat{\pi}^{(l)}_{k|g} = \frac{\sum_{i=1}^{n} w^{(l)}_{ik|g} w_{ig}}{n\hat{\pi}_g}$$

$$\hat{\mu}^{(l)}_{k|g} = \frac{\sum_{i=1}^{n} w_{ig} w^{(l)}_{ik|g} \sum_{m=1}^{M} z^{(l)}_{i,m}}{M \sum_{i=1}^{n} w_{ng} w^{(l)}_{nk|g}}$$

where $\hat{\mu}^{(l)}_{k|g} = \hat{\mu}^{(i)}_{m,k|g}$.

$$\hat{\sigma}^{(l)}_{k|g} = \frac{\sum_{i=1}^{n} w_{ig} w_{ik|g} \sum_{m=1}^{M} (z^{(l)}_{i,m} - \hat{\mu}^{(l)}_{k|g})^2}{M \sum_{i=1}^{n} w_{ig} w_{ik|g}}$$

$$\hat{\rho}^{(l)}_{k|g} = \frac{2 \sum_{i=1}^{n} w_{ig} w_{ik|g} \sum_{m=1}^{M-1} \sum_{m'=2}^{M} (z^{(l)}_{i,m} - \hat{\mu}^{(l)}_{k|g})(z^{(l)}_{i,m'} - \hat{\mu}^{(l)}_{k|g})}{\sum_{i=1}^{n} w_{ig} w_{ik|g} \left[ \sum_{m=1}^{M-1} (z^{(l)}_{i,m} - \hat{\mu}^{(l)}_{k|g})^2 + \sum_{m'=2}^{M} (z^{(l)}_{i,m'} - \hat{\mu}^{(l)}_{k|g})^2 \right]}$$

5. Compute the log-likelihood of the copula mixture model in step (4).

6. Updating the pseudo-data based on the parameter estimated in (3).

7. Repeat E-step and M-step until convergence is reached, i.e. the log-likelihood between two consecutive iterations is less than $\epsilon$ or the difference between the parameter estimates is less than a small $\epsilon$.
